# 15-PGDH Inhibition Activates the Splenic Niche to Promote Hematopoietic Regeneration

**DOI:** 10.1101/2020.09.17.302422

**Authors:** Julianne N.P. Smith, Dawn M. Dawson, Kelsey F. Christo, Alvin P. Jogasuria, Mark J. Cameron, Monika I. Antczak, Joseph M. Ready, Stanton L. Gerson, Sanford D. Markowitz, Amar B. Desai

**Author notes:** Correspondence: Amar B. Desai, Department of Medicine, Case Western Reserve University.

## Abstract

The splenic microenvironment regulates hematopoietic stem and progenitor cell (HSPC) function, particularly during demand-adapted hematopoiesis, however practical strategies to enhance splenic support of transplanted HSPCs have proven elusive. We have previously demonstrated that inhibiting 15-hydroxyprostaglandin dehydrogenase (15-PGDH), using the small molecule (+)SW033291 (PGDHi), increases bone marrow (BM) prostaglandin E2 (PGE2) levels, expands HSPC numbers, and accelerates hematologic reconstitution following BM transplantation (BMT) in mice. Here we demonstrate that the splenic microenvironment, specifically 15-PGDH high-expressing macrophages (MΦs), megakaryocytes (MKs), and mast cells (MCs), regulates steady-state hematopoiesis and potentiates recovery after BMT. Notably, PGDHi-induced neutrophil, platelet, and HSPC recovery were highly attenuated in splenectomized mice. PGDHi induced non-pathologic splenic extramedullary hematopoiesis at steady-state, and pre-transplant PGDHi enhanced the homing of transplanted cells to the spleen. 15-PGDH enzymatic activity localized specifically to MΦs, MK lineage cells, and MCs, identifying these cell types as likely coordinating the impact of PGDHi on splenic HSPCs. These findings suggest that 15-PGDH expression marks novel HSC niche cell types that regulate hematopoietic regeneration. Therefore, PGDHi provides a well-tolerated strategy to therapeutically target multiple HSC niches and to promote hematopoietic regeneration and improve clinical outcomes of BMT.

## Introduction

The spleen influences hematopoietic stem cell transplantation outcomes, yet the mechanisms regulating splenic hematopoiesis post-transplantation are not well-understood. In mice, transplanted hematopoietic stem and progenitor cells (HSPCs) home to the spleen prior to the bone marrow (BM) ^1^ and spleen-homed HSPCs demonstrate superior function relative to BM-homed HSPCs several hours post-transplant ^2^. In the days to weeks following transplantation, hematopoietic foci form in the spleen ^3,4^, corresponding to sites of HSPC proliferation and maturation. In humans, splenomegaly portends delayed neutrophil and platelet engraftment ^5^, however, splenectomy does not improve survival and has been linked to graft versus host disease and lymphoproliferative disease ^6,7^. Thus the spleen is capable of both positively and negatively regulating hematopoietic reconstitution and further investigation into the interactions between the splenic microenvironment and transplanted HSPCs is necessary.

HSPCs lodge in the spleen via CXCL12 expressed by peri-sinusoidal cells in the red pulp ^8^. In the setting of extramedullary hematopoiesis (EMH), CXCL12+ and SCF+ stromal cell populations also promote HSPC activation and myelo-erythroid progenitor cell expansion ^9^, while VCAM1+ macrophages retain HSPCs in the spleen ^10^. These findings highlight the potential therapeutic utility of strategies to promote or limit EMH, to improve hematopoietic regeneration or limit inflammation mediated by spleen-derived myeloid cells, as occurs in cardio- and neuro-vascular disease ^4,11,12^.

We have previously shown that inhibition of 15-hydroxyprostaglandin dehydrogenase (15-PGDH) expands HSPCs at steady-state, and enhances hematopoietic regeneration following transplantation and during BM failure ^13-15^. 15-PGDH inhibition (PGDHi) increases PGE2 and induces *Cxcl12* and *Scf* expression by BM stromal cells, however the impact of PGDHi on the spleen, particularly post-transplant, is not well-understood. Here we identify splenic 15-PGDH, and specifically 15-PGDH-expressing macrophages, megakaryocytes, and mast cells, as regulators of EMH and propose that targeting splenic 15-PGDH prior to transplant will enhance homing and regeneration, resulting in improved clinical outcomes.

## Results

### The spleen is critical for PGDHi-mediated hematopoietic regeneration

To determine if the spleen responds to 15-PGDH inhibition, we examined 15-PGDH expression levels in splenic tissue. Splenic 15-PGDH expression was substantially elevated versus that of BM (**Fig. 1A**). Immunohistochemical staining for 15-PGDH also revealed a striking difference in the abundance of 15-PGDH+ cells (**Fig. 1B**). While the BM displayed relatively rare 15-PGDH+ cells, which comprised smaller hematopoietic cells and megakaryocytes, splenic 15-PGDH+ cells were highly numerous, particularly within the red pulp (**Supplemental Fig. 1**). Consistent with these findings, 15-PGDH enzymatic activity was significantly higher in splenic as compared to BM cell lysates (**Fig. 1C**), demonstrating that the abundant 15-PGDH is enzymatically active. Together these results suggested that the spleen may be more sensitive than the marrow to pharmacologic 15-PGDH targeting.

**Figure 1.**
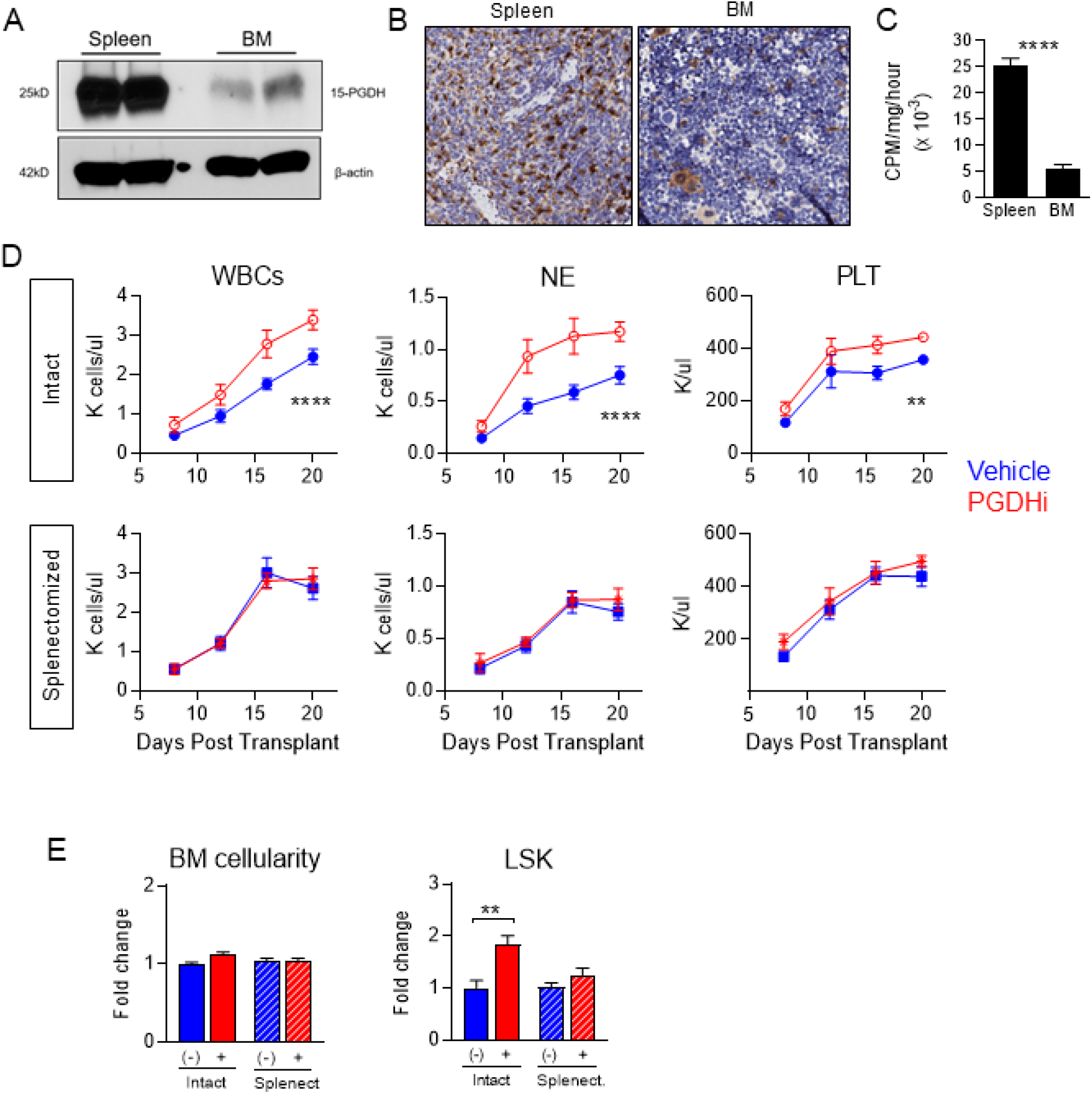
The spleen is critical for PGDHi-mediated hematopoietic regeneration. **A**. Representative detection of 15-PGDH at 25kD and β-actin at 42kD in splenocyte and bone marrow (BM) cell lysates. **B**. Representative images of 15-PGDH staining (brown) in splenic red pulp (left) and tibial BM core (right). **C**. Quantification of 15-PGDH enzymatic activity in spleen and BM, expressed as counts per minute (CPM) per mg total protein, per hour. N = 5 mice. **D**. Peripheral white blood cell (WBC), neutrophil (NE), and platelet (PLT) recovery in intact (top) and splenectomized (bottom) transplant recipients treated with either vehicle (Veh; blue) or 15-PGDH inhibitor (PGDHi; red). N = 12-15 mice/group. **E**. BM cellularity and quantification of lineage^-^ c-Kit^+^ Sca-1^+^ (LSK) cells per hindlimb of control and splenectomized recipients 20 days post-transplant, treated with Veh (-) or PGDHi (+), expressed as fold change. N = 11-14 mice per group. \*\**P* < 0.01, \*\*\*\**P* < 0.0001. Student’s t-test used for all except peripheral blood recovery, where 2-way ANOVA was used.

Having established that 15-PGDH is expressed much more highly in the spleen than the marrow, we next sought to determine whether the spleen is required for the hematopoietic protective effects of 15-PGDH inhibition (PGDHi; ^14^). To test this, we compared short-term hematologic recovery from transplant in splenectomized versus intact mice. Although vehicle-treated splenectomized mice recovered blood counts slightly faster than intact controls, as has been reported ^16,17^, splenectomy markedly attenuated the impact of PGDHi on neutrophil recovery and abrogated the impact of PGDHi on platelet recovery (**Fig. 1D**). Importantly, PGDHi-treated mice with spleens reached absolute neutrophil counts of 935 by day 12, as compared to 456 in splenectomized counterparts. Splenectomized mice also failed to show a PGDHi-dependent acceleration of total white blood cell recovery, suggesting that myeloid to lymphoid lineage skewing was not occurring. In addition, PGDHi did not enhance donor-derived HSPC numbers in the BM of splenectomized mice at day 20 (**Fig. 1E and Supplemental Fig. 2**). These data therefore establish that the spleen is required for PGDHi-mediated hematologic recovery.

**Figure 2.**
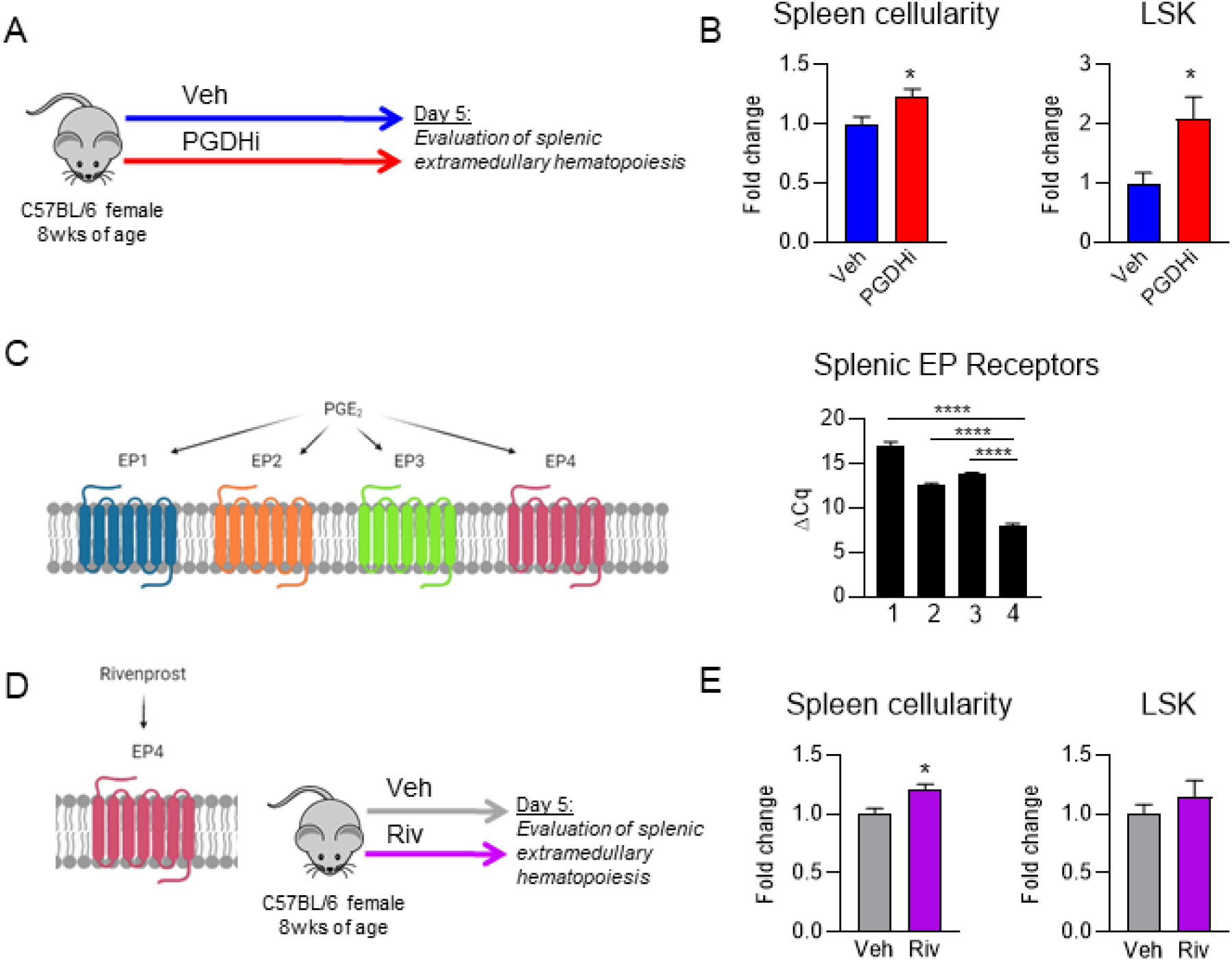
PGDHi induces splenic extramedullary hematopoiesis via EP4 activation. **A**. Schematic depicting 15-PGDH inhibition (PGDHi) in steady-state mice over the course of 5 days (9 injections). **B**. Quantification of splenic cellularity and lineage^-^ c-Kit^+^ Sca-1^+^ (LSK) cells per spleen following 5 days Veh- and PGDHi-treatment, expressed as fold change. N=12-13 mice/group for splenic cellularity and n = 7-8 mice per group for splenic LSK number. **C**. EP 1-4 (*Ptger1, 2, 3*, and *4*) expressed as delta Cq (ΔCq) relative to *B2m* control gene expression levels in CD45+ splenocytes. N = 3 mice. **D**. Schematic depcting Rivenprost administration in mice over the course of 5 days (9 doses). **E**. Quantification of splenic cellularity and LSK numbers following 5 days Veh- and Rivenprost (Riv)-treatment, expressed as fold change. N=10-11 mice/group. \**P* < 0.05, \*\*\*\**P* < 0.0001. Student’s t-test used for all except for splenic EP receptor expression, where one-way ANOVA with Tukey’s multiple comparisons test was used. PGE2 signaling diagrams created with BioRender.com.

### PGDHi induces splenic extramedullary hematopoiesis via EP4 activation

To determine if an increase in splenic EMH may underlie PGDHi-mediated hematopoietic protection post-transplant, and thus explain why splenectomized mice do not respond to PGDHi, we characterized the spleens of healthy mice treated for 5 days with PGDHi (**Fig. 2A**). PGDHi-treated mice showed significant increases in total splenic cellularity and in splenic HSPCs (**Fig. 2B**), suggesting that at homeostasis, 15-PGDH negatively regulates hematopoiesis in the spleen. PGDHi also increases BM HSPCs at homeostasis ^14^, however HSPCs are not detectable in the blood of PGDHi-treated mice (**Supplemental Fig. 3**), therefore it is unlikely that HSPC mobilization from the BM to the spleen accounts for this effect.

**Figure 3.**
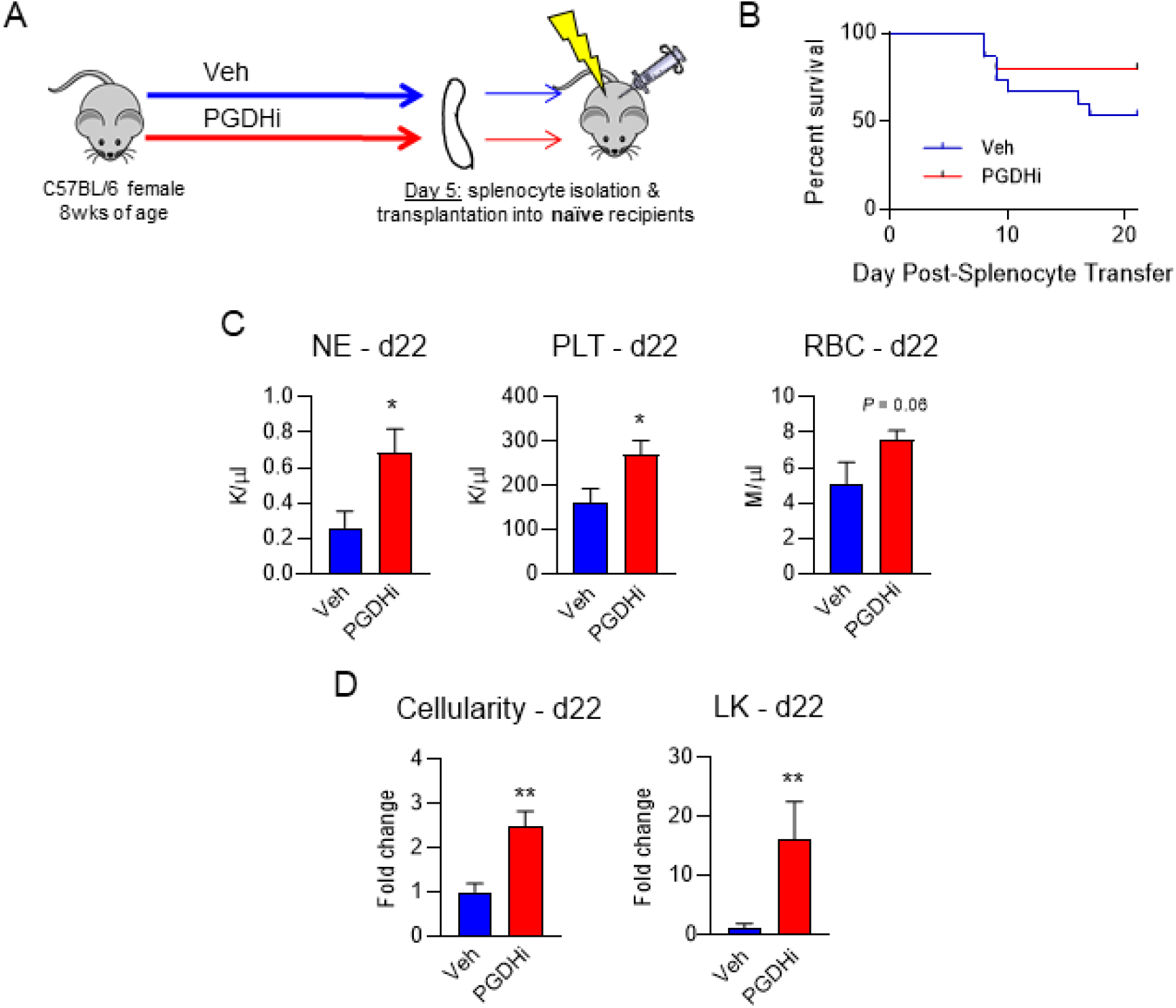
PGDHi expands functional HSPCs in the spleen. **A**. Schematic depicting the transplantation of splenocytes from PGDHi treated donors into irradiated, untreated recipients. **B**. Overall survival time of mice that received splenocytes from Veh- or PGDHi-treated donors. N=16 mice/group. Statistical testing by Log-rank (Mantel-Cox) test. **C**. Quantification of peripheral blood neutrophils (NE), platelets (PLT), and red blood cells (RBCs) in mice that received splenocytes from either Veh- or PGDHi-treated donors, 22 days post-transplant. **D**. Quantification of BM cellularity and lineage-c-Kit+ (LK) BM cells in recipient mice, 22 days post-transplant, expressed as fold change. N=6-8 mice/group. \**P* < 0.05, **P < 0.01, and ***P = 0.0008. Statistical testing of panels C-D was done by Student’s t-test, except in the case of LK cell fold change, where a Mann-Whitney test was performed.

PGE2 signals via prostaglandin receptors EP1-4 ^18^. Analysis of EP1-4 expression in splenic CD45+ cells revealed a significant predominance in the expression of the gene encoding EP4 relative to EP1, 2, and 3 (**Fig. 2C**). To determine if PGE2 signaling via EP4 may underlie the PGDHi-induced EMH, we treated mice with the EP4 specific agonist, Rivenprost ^19^ (**Fig. 2D**). Although EP4 agonism was sufficient to increase splenic cellularity, it failed to significantly expand splenic HSPCs (**Fig. 2E**). These data therefore indicate that PGDHi likely mediates splenic EMH via the actions of PGE2-EP4 signaling, but do not rule out involvement of EP1-3.

### PGDHi expands functional HSPCs in the spleen

To test whether splenic EMH corresponded to an increase in functional HSPCs in the spleen of PGDHi-treated mice, we transplanted splenocytes from PGDHi-treated donors into lethally irradiated recipients (**Fig. 3A**). A limiting cell dose of 2e6 splenocytes was chosen to assess both survival and hematologic recovery. 47% of mice that received control splenocytes succumbed to hematopoietic failure (**Fig. 3B**) evidenced by pallor, hypothermia, and lethargy (not shown). In contrast, splenocytes derived from PGDHi-treated donors conferred 20% survival. To assess hematologic recovery, surviving mice were sacrificed 22 days post-transplant. Recipients of splenocytes from PGDHi donors showed marked increases in peripheral blood neutrophils, platelets, and a trend towards increased red blood cells (**Fig. 3C**). Although the BM remained hypocellular, PGDHi donor splenocytes were associated with significantly increased engraftment of the BM in total and the lineage^-^ c-Kit^+^ immature compartment specifically (**Fig. 3D**). Together these data demonstrate that PGDHi enhances the hematopoietic capacity of the spleen to increase cellular proliferation, and expand the pool of spleen-resident HSPCs.

### Recipient PGDHi preconditioning enhances homing to the BM and splenic niches

Much like the BM, splenic hematopoiesis is regulated through the local tissue microenvironment ^9^. As PGDHi elicited splenic EMH in healthy mice, we next sought to test the therapeutic relevance of these findings and determine if pre-transplant PGDHi would increase splenic homing (**Fig. 4A**). Recipient mice treated with PGDHi prior to transplant demonstrated a 1.5-fold increase in the frequency of donor cells present in the spleen 16 hours post-transplant (**Fig. 4B**). Pre-transplant PGDHi also increased the homing of transplanted cells to the BM (**Fig. 4C**), suggesting that 15-PGDH inhibition enhances the capacity of both the splenic and the BM microenvironment to recruit and support engrafting cells.

**Figure 4.**
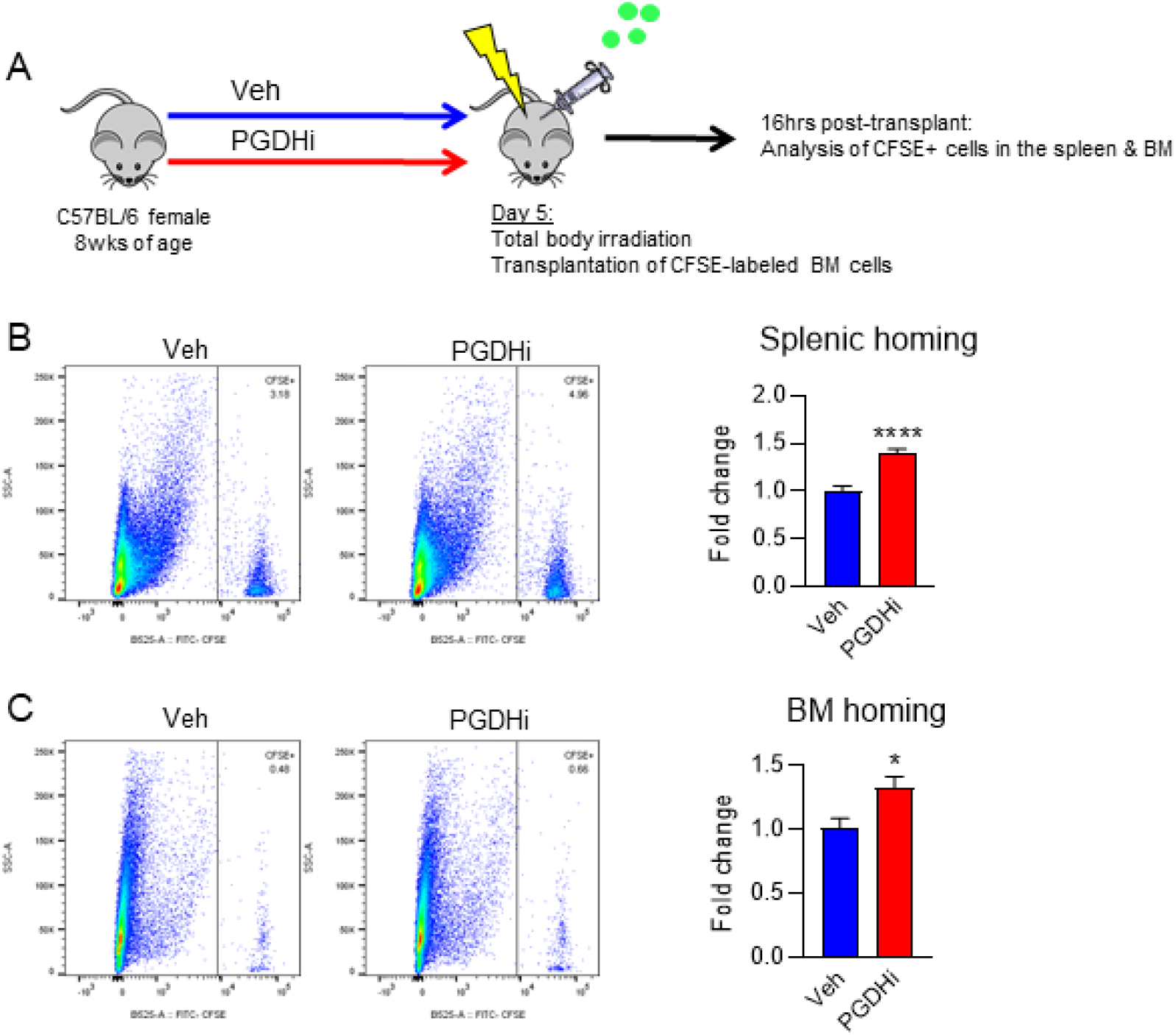
Recipient PGDHi preconditioning enhances homing to the BM and splenic niches. **A**. Schematic depicting the analysis of homing into PGDHi-pretreated recipients. **B**. Representative flow cytometry plots depicting the detection of CFSE+ cells among total splenocytes isolated 16 hours post-transplantation of pretreated mice. Graph represents fold change in the frequency of CFSE+ splenocytes. **C**. Representative flow cytometry plots depicting the detection of CFSE+ cells among total bone marrow (BM) cells isolated 16 hours post-transplant in Veh- and PGDHi-pretreated mice. Graph represents fold change in the frequency of CFSE+ BM cells. N = 11-13 mice/group. *P < 0.05, ****P < 0.0001. Statistical testing by Student’s t-test.

### PGDHi elicits a pro-hematopoietic gene expression signature in the spleen

We next sought to identify whether these PGDHi-mediated effects were associated with a pro-hematopoietic gene expression signature in the spleen and BM. As 15-PGDH expression localized to the splenic red pulp, we analyzed the expression of a number of hematopoietic niche-related genes ^20-22^ **(Fig. 5A)** in lymphoid-depleted BM and splenic cells following 5 days of vehicle or PGDHi treatment. Consistent with our findings that PGDHi induces splenic EMH and promotes homing to the splenic and BM niches, we found that a number of factors were modestly induced including the niche retentive factors *Spp1 and Vcam1* ^*10,23*^, and the quiescence-promoting factor *Kitl* ^24^ **(Fig. 5B)**. PGDHi elicited moderate induction of the *Ackr1* gene, which has been implicated in maintaining hematopoietic quiescence via the macrophage niche ^25^, specifically in the spleen (**Supplemental Fig. 4**). The sum of the individual gene expression changes across the panel of hematopoietic niche-associated factors revealed a significant increase in both organs, however, suggesting that PGDHi induces a pro-niche response that facilitates post HST engraftment.

**Figure 5.**
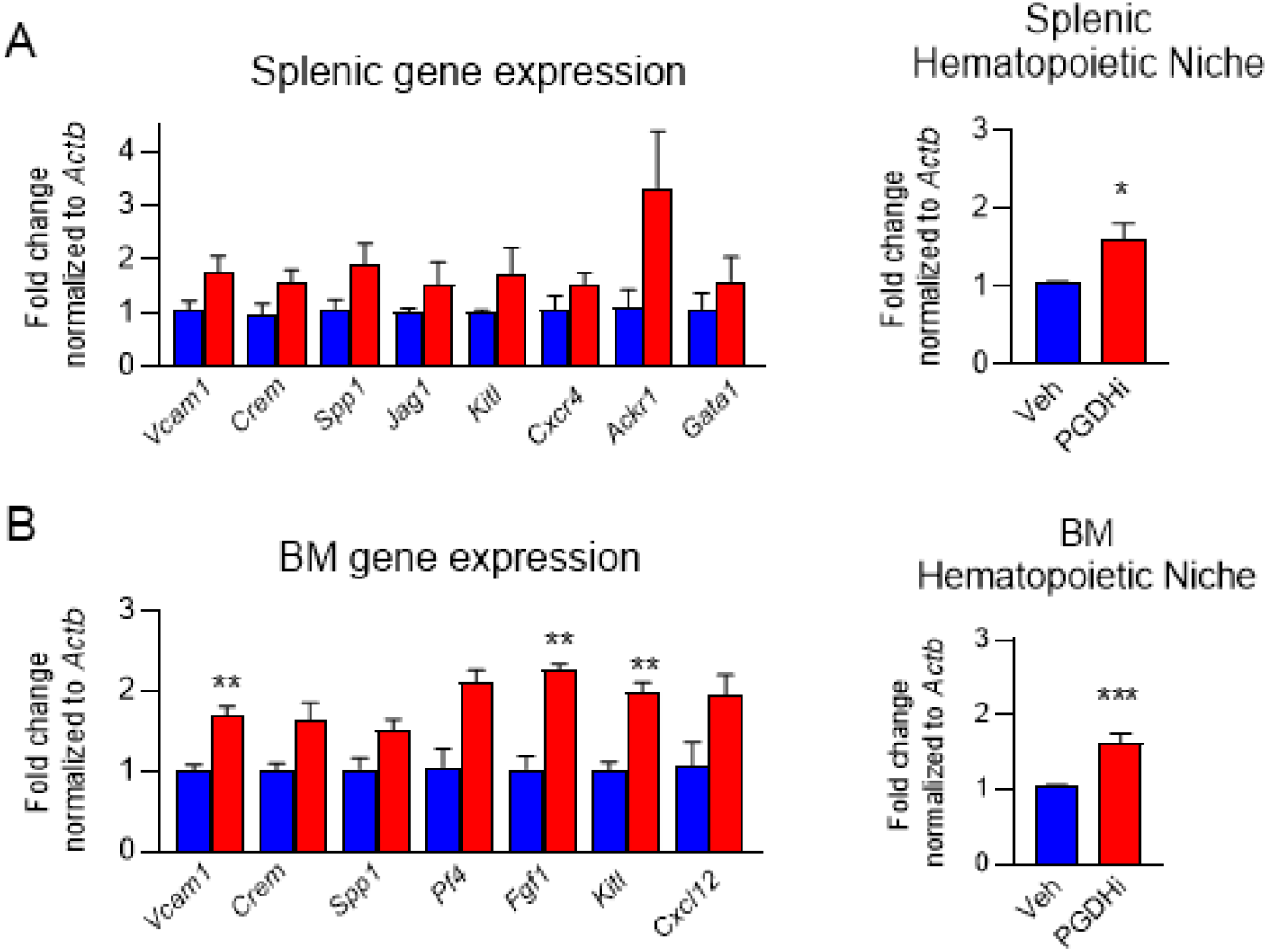
PGDHi elicits a pro-hematopoietic gene expression signature in the spleen and BM. **A**. Hematopoietic niche-related genes assayed. **B**. Relative expression of indicated genes in lymphoid-depleted splenocytes from vehicle-(blue) and PGDHi-(red) treated mice, normalized to *Actb* (left). Fold change in the expression of all hematopoietic niche-related genes, listed in panel A (right). **C**. Relative expression of indicated genes in lymphoid-depleted bone marrow (BM) cells from vehicle-(blue) and PGDHi-(red) treated mice, normalized to *Actb* (left). Fold change in the expression of all hematopoietic niche-related genes, listed in panel A (right). N = 3 mice/group. *P = 0.02, **P < 0.005, ***P = 0.0001. Statistical testing by Student’s t-test.

### 15-PGDH is highly enriched in splenic and BM mast cells, megakaryocytes, and macrophages

Given the functional significance of splenic 15-PGDH, we sought to identify the cellular sources of 15-PGDH activity in murine spleen. Immunohistochemical 15-PGDH staining principally identified hematopoietic cell types, including megakaryocytes; whereas, splenic stroma revealed very low 15-PGDH activity (data not shown). Isolation of bulk CD45+ hematopoietic cells showed no enrichment of 15-PGDH enzymatic activity per milligram protein compared to total unfractionated splenocytes (**Fig. 6A**). To determine whether immature hematopoietic cells are 15-PGDH+, and thus direct targets of PGDHi, we compared hematopoietic lineage negative and lineage positive cells. 15-PGDH activity per milligram protein was very low in the immature, as compared to the mature cell fraction, suggesting that 15-PGDH localizes to a subset of mature cells. Analysis of myeloid cells by CD11b fractionation demonstrated relative enrichment (data not shown), thus we reasoned that the major cellular sources of 15-PGDH include a myeloid cell type. As prostanoid signaling is known to regulate macrophages (MΦ), megakaryocytes (MK), and mast cells (MC), we measured activity specifically within these fractions. F4/80+ MΦs accounted for the highest level of enzyme activity per milligram protein, but significant enrichment was also measured in CD61+ MKs, and FcεR1a+ MCs. Among these cell types, F4/80+ cells are the most numerous in the spleen, accounting for 19.3% of nucleated splenocytes (**Fig. 6B**). In contrast, CD61+ and FcεR1a+ cells represented 7.6% and 0.3% of nucleated splenocytes, respectively. As a reference, CD3+ cells represented 28.2% of cells analyzed (data not shown). Notably, 15-PGDH activity also localized to CD61+, FcεR1a+, and F4/80+ cells in the BM (**Fig. 6C**), though these levels were much lower than those of the corresponding splenic populations. Among 15-PGDH+ cell types, F4/80+ cells were also the most numerous in the BM (**Fig. 6D**). These data therefore implicate splenic MΦs as the predominant cellular targets of PGDHi treatment in hematopoietic tissue.

**Figure 6.**
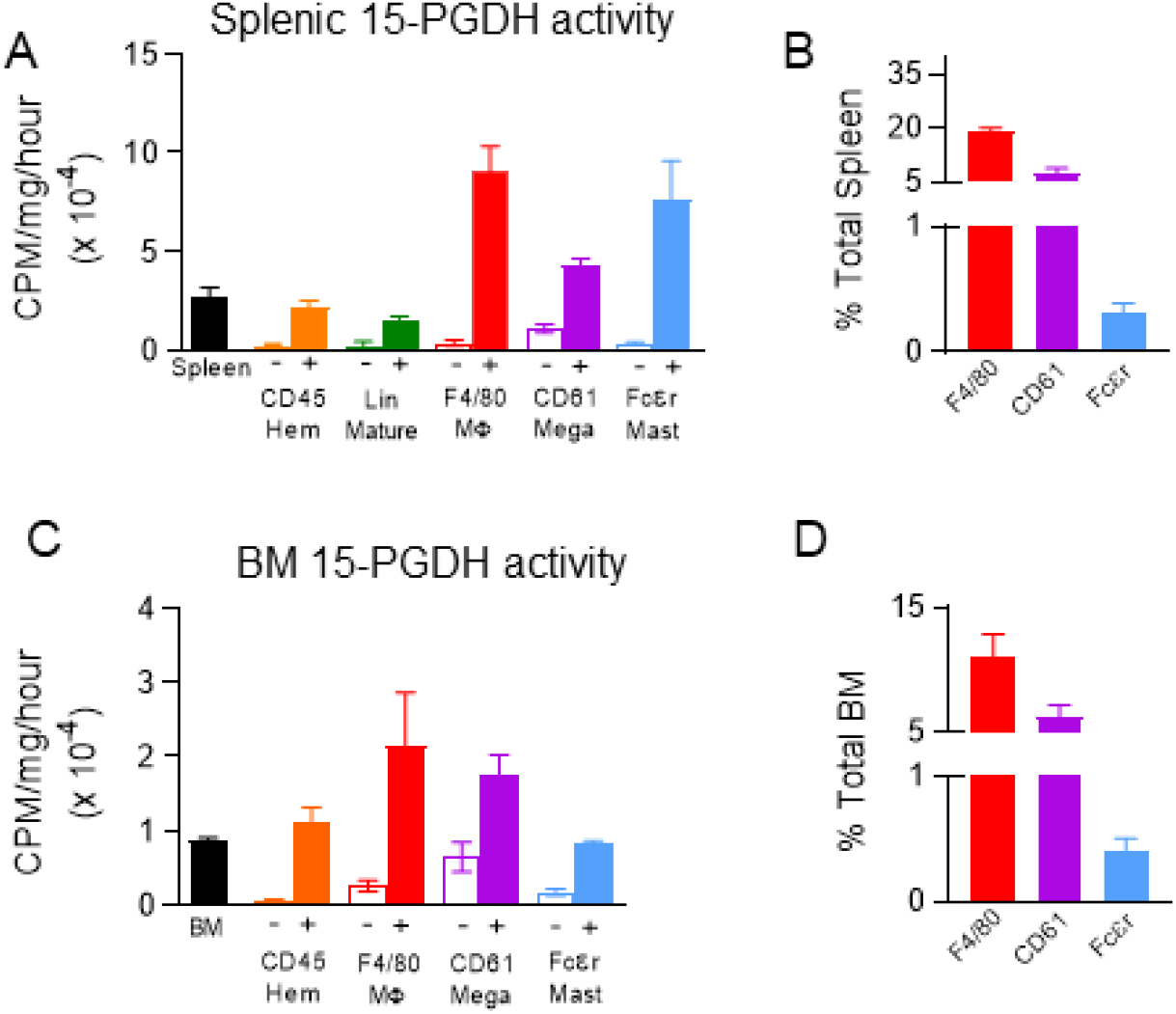
15-PGDH activity is highly enriched in splenic and marrow mast cells, megakaryocytes, and macrophages. **A**. Quantification of 15-PGDH enzymatic activity in total unfractionated spleen and in splenic CD45, hematopoietic Lineage (Lin), F4/80, CD61, and Fcεr1a -/+ fractions, respectively, expressed as CPM per mg total protein, per hour. Open bars indicate each marker’s negative population. N = 3-7 mice/cell population. **B**. Quantification of the frequency of F4/80+, CD61+, and Fcεr1a+ cells in the murine spleen. N = 3 mice. **C**. Quantification of 15-PGDH enzymatic activity in total unfractionated bone marrow (BM) and in BM CD45, F4/80, CD61, and Fcεr1a -/+ fractions, respectively, expressed as counts per minute (CPM) per mg total protein, per hour. Open bars indicate each marker’s negative population. N = 3-7 mice/cell population. **D**. Quantification of the frequency of F4/80+, CD61+, and Fcεr1a+ cells in the murine BM. N = 3 mice.

### 15-PGDH localization and enzymatic activity is conserved in human BM

To determine if 15-PGDH expression patterns are conserved between murine and human hematopoietic tissue, we evaluated healthy human biopsies and aspirates. 15-PGDH+ marrow cells were readily detectable in all human biopsies examined (**Fig. 7A**). Positive cells varied in size and morphology, and included pyramidal, elongated, and round cells. To identify the cellular localization of 15-PGDH activity, we separated cells from human BM aspirates on the basis of surface marker expression. Relative to the activity of total BM, FcεR1a+ MCs demonstrated 115-fold higher levels of specific 15-PGDH activity (**Fig. 7B**). CD14+ MΦs and CD61+ MK-lineage cells also demonstrated enzyme activity enrichment, though to lesser degrees than FcεR1a+ cells. These results establish that FcεR1a+ MCs, CD61+ MKs, and CD14+ MΦs are robust sources of 15-PGDH activity in human marrow, and thus may comprise a novel and therapeutically-targetable human HSC niche.

**Figure 7.**
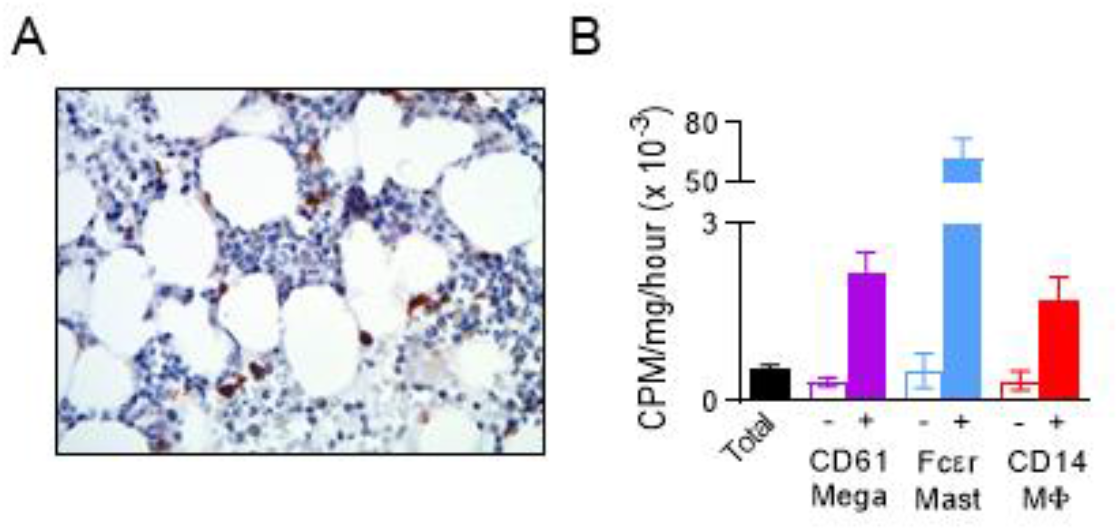
15-PGDH localization and enzymatic activity is conserved in human BM. **A**. Representative image of 15-PGDH staining (brown) in a human bone marrow (BM) biopsy. **B**. Quantification of 15-PGDH enzymatic activity in total (unfractionated) human BM as compared to CD61, Fcεr1a, and CD14 -/+ fractions, expressed as CPM per mg total protein, per hour. Open bars indicate each marker’s negative population. N = 3-5 donors.

## Discussion

Previously, we established that 15-PGDH regulates hematopoietic, colonic epithelial, and hepatic tissue regeneration ^13-15^. Pharmacologic 15-PGDH inhibition or loss of *Hpgd* expression (that encodes 15-PGDH) elevated BM prostaglandin E2, D2 and F2a levels, increased peripheral neutrophils, and expanded the BM HSPC compartment. PGDHi also enhanced the progenitor activity, homing, and reconstituting ability of murine BM, human BM, and umbilical cord blood. PGDHi induced the expression of niche factors in the BM. However, because splenic colony forming units and splenic HSPCs were also increased, we hypothesized that splenic 15-PGDH also negatively regulates hematopoietic regeneration. Here, we demonstrate robust 15-PGDH expression in splenic red pulp, which localizes to MΦs, MKs, and MCs. Our observation that the spleen is required for PGDHi responses post-transplant, advances current understanding of splenic EMH and identifies potential therapeutic targets within the splenic HSPC niche.

Prostaglandin E2 is an arachidonic acid derivative capable of increasing HSPC numbers *in vivo* and *in vitro* ^26-28^. Although PGE2 can be produced by a multitude of cells types, osteoblasts, endothelial cells, and monocyte-macrophage lineage cells have been most extensively characterized in the HSPC microenvironment (reviewed in ^29^). HSPCs and non-hematopoietic microenvironmental cell types express PGE2 receptors ^30^, and agonism of EP2 and EP4 receptors on HSPCs activates Wnt signaling and increases the expression of anti-apoptotic and pro-proliferative gene programs ^31^. PGE2 stimulation also increases HSPC CXCR4 expression ^32^, thus enhancing homing capacity. Recently, a role for MΦ-derived PGE2 in facilitating erythropoietin-induced erythropoiesis was reported by Chen *et al*. ^33^. Clinical trials have evaluated the *ex vivo* stimulation of human cord blood with the long-acting PGE2 analog dimethyl-PGE2 (dmPGE2) as a strategy to enhance engraftment ^34^. *Ex vivo* stimulation avoids the potential for off-target dmPGE2-induced toxicity, however, our data together with the radioprotective and erythropoiesis-promoting effects of PGE2 ^26,27,33^, demonstrate the potential for additional benefit from strategies to elevate tissue PGE2 levels in transplant recipients. Future studies to evaluate dmPGE2 *ex vivo* graft stimulation combined with pre-transplant PGDHi recipient conditioning are therefore warranted.

In the adult, hematopoiesis takes place primarily in the BM where HSPCs are regulated by perivascular stromal cells, endothelial cells, MΦs, and MKs ^35-39^. The red pulp of the spleen serves as an alternative HSPC microenvironment when the BM is dysfunctional, however (reviewed in ^40^), and provides myelopoiesis and erythropoiesis in response to infection ^41^, inflammation ^42^, and physical and psychological stress ^43,44^. Rare HSPCs are found in murine spleen under homeostatic conditions ^9^, however, and recent reports demonstrate human splenic EMH in the absence of disease ^45^. Here we establish that PGDHi induces nonpathologic splenic EMH. Splenic endothelial and Tcf21+ stromal cells were recently shown to regulate EMH and particularly myeloerythroid lineage differentiation ^9^. Our studies do not directly address the impact of PGDHi on spleen stroma, but 15-PGDH activity was very low in splenic CD45-cells, indicating that stromal cells are not likely direct PGDHi targets.

PGDHi likely increases splenic cellularity via PGE2 stimulation of EP4 receptor, as EP4 specific agonism recapitulates this phenotype. EP4 agonism is not sufficient to expand the pool of splenic HSPCs, however, suggesting activation of additional EP receptors is required for the induction of splenic EMH by PGDHi. Moreover, PGDHi potentiates splenic homing of transplanted cells. As the spleen is associated with delayed engraftment in some transplant patients ^5^, our data suggest that PGDHi may provide an alternative to splenectomy or splenic irradiation. Whether PGDHi improves hematopoietic function in other pathophysiologic states that involve splenic EMH, such as infection or blood loss, is an intriguing question.

BM MKs enforce HSC quiescence via CXCL4 and TGFβ but take on an FGF1-dependent HSC activating role upon hematologic stress ^38,39^. Similarly, MΦs maintain quiescence and niche retention at steady-state in part through activities of the atypical chemokine receptor 1 (*Ackr1*; ^25^), and VCAM1 (^10^ and reviewed in ^46^), but exacerbate inflammation and regulate HSPC differentiation in pathologic conditions ^47^. Our finding that splenic MKs and MΦs express high levels of enzymatically-active 15-PGDH suggests that these cell types participate in the PGDHi-dependent regulation of splenic EMH. PGE2 limits inflammation in some contexts ^48^, and irradiation potentiates the inflammatory state of MΦs ^49^, thus it is possible that PGDHi attenuates MΦ activation to preserve splenic niche function. Additionally, PGE2 inhibits TGFβ signaling ^50^, therefore, PGDHi treatment may modulate the role of splenic MKs from promoting HSC quiescence to activation. Our work also implicates MCs as components of the splenic EMH microenvironment. MCs are rich in histamine- and leukotriene-containing granules however, and thus are poised to rapidly regulate the local tissue microenvironment. Moreover, leukotriene B4 has been hypothesized to promote HSPC differentiation at the expense of self-renewal ^29^, and PGE2 suppresses MC degranulation in anaphylaxis ^51^. Increased splenic myelopoiesis and thrombopoiesis have also been observed in MC deficient mice ^52^. Future studies to evaluate the impact of PGDHi specifically on splenic MCs, MKs, and MΦs are warranted.

In conclusion, 15-PGDH is highly expressed and enzymatically-active in the murine spleen. Pre-transplant PGDHi induces a pro-niche gene signature in the splenic and BM microenvironments and induces splenic EMH, which translates to an increase in the homing of transplanted cells to the spleen. We find that the spleen is required for PGDHi-mediated leukocyte and platelet reconstitution and BM HSPC engraftment. This likely owes to a network of 15-PGDH+ macrophages, megakaryocytes, and rare mast cells in the spleen. Therefore, our work identifies a novel pharmacologic strategy and the corresponding cellular targets that regulate extramedullary hematopoiesis. Small molecule 15-PGDH inhibition represents a novel therapeutic strategy to utilize the splenic microenvironment post-transplant and likely in other disease states where rapid hematopoietic regeneration is needed.

## Methods

### Reagents

15-PGDH inhibitors (+)SW033291 and (+)SW209415 were previously described ^13,14^, and provided by Dr. Sanford Markowitz. (+)SW033291 was prepared in a vehicle of 10% ethanol, 5% Cremophor EL, 85% dextrose-5 water, at 125ug/200ul for use at 5mg/kg for a 25g mouse, and administered by intraperitoneal (I.P.) injection, twice per day, 6-8 hours apart. (+)SW209415 was prepared as previously described ^13^, and administered at 2.5mg/kg I.P., twice per day. Rivenprost (Cayman Chemical) was prepared in a vehicle of PBS for use at 30ug/kg and administered by I.P. injection, twice per day, 6-8 hours apart. Carboxyfluorescein succinimidyl ester (CFSE) Cell Trace was purchased from Invitrogen.

### Animals

Animals were housed in the AAALAC accredited facilities of the CWRU School of Medicine. Husbandry and experimental procedures were approved by the Case Western Reserve University Institutional Animal Care and Use Committee (IACUC) in accordance with approved IACUC protocols 2013-0182 and 2019-0065. Steady-state and transplantation analyses were performed on 8wk old female C57BL/6J mice obtained from Jackson Laboratories. B6.SJL-Ptprca Pepcb/BoyJ and splenectomized C57BL/6 mice were obtained from Jackson Laboratories. All animals were observed daily for signs of illness. Mice were housed in standard microisolator cages and maintained on a defined, irradiated diet and autoclaved water.

### Western Blotting

Cells were lysed using RIPA lysis buffer containing protease inhibitors. Lysates were centrifuged 10,000 rpm for 10 minutes at 4C. Protein concentrations were determined by BCA assay. Proteins were separated using 4-12% SDS-PAGE gels, then transferred to PVDF membranes, and probed with antibodies recognizing murine 15-PGDH (kindly provided by Dr. Sanford Markowitz), and β–actin (Sigma, A5441).

### Histological and Immunohistochemical Analysis

Animals were harvested via CO2 inhalation followed by cervical dislocation. Whole spleens or tibial bone marrow plugs were fixed in 10% neutral buffered formalin. Samples were transferred to PBS and shipped to HistoWiz where they were embedded in paraffin, and sectioned at 4μm. Immunohistochemistry was performed according to Histowiz protocols (https://home.histowiz.com/faq/). Histowiz defines their standard methods as the use of a Bond Rx autostainer (Leica Biosystems) with enzyme treatment using standard protocols, and detection via Bond Polymer Refine Detection (Leica Biosystems) according to manufacturer’s protocol. Whole slide scanning (40x) was performed on an Aperio AT2 (Leica Biosystems).

### Measurement of 15-PGDH Enzymatic Activity

Splenic lysates were prepared using the Precellys 24 homogenizer, in a lysis buffer containing 50mM Tris HCl, 0.1mM DTT, and 0.1mM EDTA. Bone marrow was flushed, pelleted, and lysed using the same buffer, with sonication. Enzymatic activity was measured by following the transfer of tritium from a tritiated PGE2 substrate to glutamate by coupling 15-PGDH to glutamate dehydrogenase ^53^. Activity was expressed as counts per minute, per mg total protein assayed.

### Bone Marrow Transplantation

Mice were exposed to 10Gy total body irradiation from a cesium source, followed immediately by administration of PGDHi or vehicle control. 16-18hrs later, mice received 1e6 whole bone marrow cells by retroorbital injection, followed immediately by a second I.P. administration of PGDHi or vehicle control. Recipients continued to receive twice daily I.P. injections of PGDHi or vehicle.

### Complete Blood Count Analysis

Peripheral blood was collected into Microtainer EDTA tubes (Becton-Dickinson) by submandibular cheek puncture. Blood counts were analyzed using a Hemavet 950 FS hematology analyzer.

### Quantification of HSPCs and Splenic Cell Types

Bone marrow cells were obtained by flushing hindlimb bones and splenocytes were obtained by mincing spleens. Cellularity was measured following red blood cell lysis. Cells were stained with antibodies against CD45R/B220 (RA3-6B2), CD11b (M1/70), CD3e (500A2), Ly-6G and Ly6C (RB6-8C5), TER-119 (TER-119), Ly-6A/E (D7), CD117 (2B8), F4/80 (Cl:A3-1), CD61 (2C9.G2), and Fcer1 alpha (MAR-1), and data was acquired on an LSRII flow cytometer (BD Biosciences). Analysis was performed on FlowJo software (TreeStar).

### Cell Separation

Single cell suspensions were generated from spleen and marrows. Cells were isolated by surface marker expression using Miltenyi microbead kits and LS column separation according to manufacturer’s instructions. 15-PGDH enzymatic activity was measured in cell fractions, or in unfractionated splenocytes or marrow cells, as described above and previously reported ^14^.

### RNA Extraction and Quantitative PCR

CD45+ splenocytes or CD3e- and B220-depleted bone marrow cells and splenocytes were isolated, as described above. Cells were lysed and RNA extracted using the RNeasy MiniKit (QIAGEN) with on-column DNase treatment, according to the manufacturer’s protocol. cDNA was synthesized using the PrimeScript RT Reagent Kit (Takara) following manufacturer’s instructions. Real time PCR measurement was performed in a 20ul reaction containing 1ul cDNA template and a 1:20 dilution of primer/probe with 1X Accuris Taq DNA polymerase. Samples were run on a CFX96 optical module (Bio-Rad). Thermal cycling conditions were 95C for 3 minutes, followed by 50 cycles of 95C for 15 seconds and 60C for 1 minute. Murine probe/primer sets for all genes assayed were obtained from Life Technologies and were as follows: *B2m* Mm00437762_m1, *Ptger1* Mm00443098_g1, *Ptger2* Mm00436051_m1, *Ptger3* Mm01316856_m1, *Ptger4* Mm00436053_m1, *Actb* Mm02619580_g1, *Vcam1* Mm01320970_m1, *Crem* Mm04336053_g1, *Spp1* Mm00436767_m1, *Jag1* Mm00496902_m1, *Kitl* Mm00442972_m1, *Cxcr4* Mm01996749_s1, *Ackr1* Mm00515642_g1, *Gata1* Mm01352636_m1, *Pf4* Mm00451315_g1, *Fgf1* Mm00438906_m1, and *Cxcl12* Mm00445553_m1. For each reverse transcription reaction, Cq values were determined as the average values obtained from three independent real-time PCR reactions.

### Splenocyte Transplantation

Donor mice were treated for five days with PGDHi or vehicle control, twice daily, by I.P. injection. Two hours following the ninth administration, mice were sacrificed and spleens were dissected and a single cell suspension was generated. Recipient mice were conditioned with 10Gy irradiation 20hrs prior to the transplantation of 2e6 splenocytes by retroorbital injection.

### Bone Marrow Homing Analysis

Bone marrow was labeled with 5μM CellTrace CFSE and 10e6 cells were transplanted into recipient mice that had been treated for 5 days with PGDHi or vehicle control, and conditioned with 10Gy total body irradiation 12 hours prior to transplant. 16 hours post-transplant, mice were sacrificed and CFSE+ cells were quantified in the spleen and bone marrow flow cytometrically.

### Human Tissue Procurement

De-identified adult bone marrow aspirates were obtained from the CWRU Hematopoietic Biorepository with permission from the Institutional Review Board. Human bone marrow aspirates were depleted of red blood cells prior to cell fractionation.

### Statistical Analysis

All values were tabulated graphically with error bars corresponding to standard error of the means. Analysis was performed using GraphPad Prism software. Unpaired two-tailed Student’s t-test was used to compare groups, unless otherwise noted. For peripheral blood recovery kinetic analysis, 2-way ANOVA was used to test the effect of drug treatment.

## Authorship Contributions

Julianne N.P. Smith: Conception and Design, Collection of Data, Data Interpretation, Manuscript Writing

Dawn M. Dawson: Conception and Design, Data Interpretation

Kelsey F. Christo: Collection of Data

Alvin P. Jogasuria: Collection of Data

Mark J. Cameron: Conception and Design

Monika I. Antczak: Chemical Compound Purification and Quality Control

Joseph M. Ready: Chemical Compound Purification and Quality Control

Stanton L. Gerson: Conception and Design, Data Interpretation

Sanford D. Markowitz: Conception and Design, Data Interpretation, Manuscript Writing

Amar B. Desai: Conception and Design, Data Interpretation, Manuscript Writing, Final Approval of Manuscript

## Acknowledgments

This work was supported by NIH grants R35 CA197442, K99 HL135740, and T32 EB005583, and by the Radiation Resources Core Facility (P30CA043703), the Hematopoietic Biorepository and Cellular Therapy Core Facility (P30CA043703), the Tissue Resources Core Facility (P30CA043703), and the Cytometry & Imaging Microscopy Core Facility of the Case Comprehensive Cancer Center (P30CA043703).

---

**Table 1.** Hematopoietic niche-related genes assayed.

| Gene symbol | Gene name |
| --- | --- |
| Vcam1 | Vascular cell adhesion molecule 1 |
| Crem | cAMP responsive element modulator |
| Spp1 | Osteopontin |
| Jag1 | Jagged 1 |
| Kitl | Kit ligand |
| Cxcr4 | CXCR4 |
| Ackr1 | Atypical chemokine receptor 1 (Duffy blood group) |
| Gata1 | GATA binding protein 1 |
| Pf4 | Platelet factor 4 |
| Fgf1 | Fibroblast growth factor 1 |
| Cxcl12 | CXCL12 |

## References

1. Kollet O, Spiegel A, Peled A, et al. Rapid and efficient homing of human CD34(+)CD38(-/low)CXCR4(+) stem and progenitor cells to the bone marrow and spleen of NOD/SCID and NOD/SCID/B2m(null) mice. Blood. 2001;97(10):3283–3291.

2. Szilvassy SJ, Bass MJ, Van Zant G, Grimes B. Organ-selective homing defines engraftment kinetics of murine hematopoietic stem cells and is compromised by Ex vivo expansion. Blood. 1999;93(5):1557–1566.

3. Cao YA, Wagers AJ, Beilhack A, et al. Shifting foci of hematopoiesis during reconstitution from single stem cells. Proc Natl Acad Sci U S A. 2004;101(1):221–226.

4. Short C, Lim HK, Tan J, O’Neill HC. Targeting the Spleen as an Alternative Site for Hematopoiesis. Bioessays. 2019;41(5):e1800234.

5. Akpek G, Pasquini MC, Logan B, et al. Effects of spleen status on early outcomes after hematopoietic cell transplantation. Bone Marrow Transplant. 2013;48(6):825–831.

6. Ringden O, Nilsson B. Death by graft-versus-host disease associated with HLA mismatch, high recipient age, low marrow cell dose, and splenectomy. Transplantation. 1985;40(1):39–44.

7. Sundin M, Le Blanc K, Ringden O, et al. The role of HLA mismatch, splenectomy and recipient Epstein-Barr virus seronegativity as risk factors in post-transplant lymphoproliferative disorder following allogeneic hematopoietic stem cell transplantation. Haematologica. 2006;91(8):1059–1067.

8. Miwa Y, Hayashi T, Suzuki S, et al. Up-regulated expression of CXCL12 in human spleens with extramedullary haematopoiesis. Pathology. 2013;45(4):408–416.

9. Inra CN, Zhou BO, Acar M, et al. A perisinusoidal niche for extramedullary haematopoiesis in the spleen. Nature. 2015;527(7579):466–471.

10. Dutta P, Hoyer FF, Grigoryeva LS, et al. Macrophages retain hematopoietic stem cells in the spleen via VCAM-1. J Exp Med. 2015;212(4):497–512.

11. Nahrendorf M, Dutta P. Targeting splenic hematopoietic stem cells in cardiovascular disease. Oncotarget. 2015;6(24):19918–19919.

12. Wang Z, He D, Zeng YY, et al. The spleen may be an important target of stem cell therapy for stroke. J Neuroinflammation. 2019;16(1):20.

13. Desai A, Zhang Y, Park Y, et al. A second-generation 15-PGDH inhibitor promotes bone marrow transplant recovery independently of age, transplant dose and granulocyte colony-stimulating factor support. Haematologica. 2018;103(6):1054–1064.

14. Zhang Y, Desai A, Yang SY, et al. TISSUE REGENERATION. Inhibition of the prostaglandin-degrading enzyme 15-PGDH potentiates tissue regeneration. Science. 2015;348(6240):aaa2340.

15. Smith JNP, Otegbeye F, Jogasuria AP, et al. Inhibition of 15-PGDH Protects Mice from Immune-mediated Bone Marrow Failure. Biol Blood Marrow Transplant. 2020.

16. Smith LH, McKinley TW, Jr. Recovery from radiation injury with and without bone marrow transplantation: effects of splenectomy. Radiat Res. 1970;44(1):248–261.

17. Tanaka N. Experimental studies on role of spleen in recovery from radiation injury in mice. II. Effect of splenectomy on hematological findings and iron metabolism in mice following x-irradiation. Hiroshima J Med Sci. 1966;15(4):325–346.

18. Markovic T, Jakopin Z, Dolenc MS, Mlinaric-Rascan I. Structural features of subtype-selective EP receptor modulators. Drug Discov Today. 2017;22(1):57–71.

19. Ninomiya T, Hosoya A, Hiraga T, et al. Prostaglandin E(2) receptor EP(4)-selective agonist (ONO-4819) increases bone formation by modulating mesenchymal cell differentiation. Eur J Pharmacol. 2011;650(1):396–402.

20. Pinho S, Frenette PS. Haematopoietic stem cell activity and interactions with the niche. Nat Rev Mol Cell Biol. 2019;20(5):303–320.

21. Maupin KA, Himes ER, Plett AP, et al. Aging negatively impacts the ability of megakaryocytes to stimulate osteoblast proliferation and bone mass. Bone. 2019;127:452–459.

22. Saxena S, Ronn RE, Guibentif C, Moraghebi R, Woods NB. Cyclic AMP Signaling through Epac Axis Modulates Human Hemogenic Endothelium and Enhances Hematopoietic Cell Generation. Stem Cell Reports. 2016;6(5):692–703.

23. Stier S, Ko Y, Forkert R, et al. Osteopontin is a hematopoietic stem cell niche component that negatively regulates stem cell pool size. J Exp Med. 2005;201(11):1781–1791.

24. Oguro H, Ding L, Morrison SJ. SLAM family markers resolve functionally distinct subpopulations of hematopoietic stem cells and multipotent progenitors. Cell Stem Cell. 2013;13(1):102–116.

25. Hur J, Choi JI, Lee H, et al. CD82/KAI1 Maintains the Dormancy of Long-Term Hematopoietic Stem Cells through Interaction with DARC-Expressing Macrophages. Cell Stem Cell. 2016;18(4):508–521.

26. Porter RL, Georger MA, Bromberg O, et al. Prostaglandin E2 increases hematopoietic stem cell survival and accelerates hematopoietic recovery after radiation injury. Stem Cells. 2013;31(2):372–383.

27. Hoggatt J, Singh P, Stilger KN, et al. Recovery from hematopoietic injury by modulating prostaglandin E(2) signaling post-irradiation. Blood Cells Mol Dis. 2013;50(3):147–153.

28. Cutler C, Multani P, Robbins D, et al. Prostaglandin-modulated umbilical cord blood hematopoietic stem cell transplantation. Blood. 2013;122(17):3074–3081.

29. Hoggatt J, Pelus LM. Eicosanoid regulation of hematopoiesis and hematopoietic stem and progenitor trafficking. Leukemia. 2010;24(12):1993–2002.

30. Frisch BJ, Porter RL, Gigliotti BJ, et al. In vivo prostaglandin E2 treatment alters the bone marrow microenvironment and preferentially expands short-term hematopoietic stem cells. Blood. 2009;114(19):4054–4063.

31. Goessling W, North TE, Loewer S, et al. Genetic interaction of PGE2 and Wnt signaling regulates developmental specification of stem cells and regeneration. Cell. 2009;136(6):1136–1147.

32. Hoggatt J, Singh P, Sampath J, Pelus LM. Prostaglandin E2 enhances hematopoietic stem cell homing, survival, and proliferation. Blood. 2009;113(22):5444–5455.

33. Chen Y, Xiang J, Qian F, et al. Epo-receptor signaling in macrophages alters the splenic niche to promote erythroid differentiation. Blood. 2020.

34. Maung KK, Horwitz ME. Current and future perspectives on allogeneic transplantation using ex vivo expansion or manipulation of umbilical cord blood cells. Int J Hematol. 2019;110(1):50–58.

35. Ding L, Saunders TL, Enikolopov G, Morrison SJ. Endothelial and perivascular cells maintain haematopoietic stem cells. Nature. 2012;481(7382):457–462.

36. Boettcher S, Gerosa RC, Radpour R, et al. Endothelial cells translate pathogen signals into G-CSF-driven emergency granulopoiesis. Blood. 2014;124(9):1393–1403.

37. Ludin A, Itkin T, Gur-Cohen S, et al. Monocytes-macrophages that express alpha-smooth muscle actin preserve primitive hematopoietic cells in the bone marrow. Nat Immunol. 2012;13(11):1072–1082.

38. Bruns I, Lucas D, Pinho S, et al. Megakaryocytes regulate hematopoietic stem cell quiescence through CXCL4 secretion. Nat Med. 2014;20(11):1315–1320.

39. Zhao M, Perry JM, Marshall H, et al. Megakaryocytes maintain homeostatic quiescence and promote post-injury regeneration of hematopoietic stem cells. Nat Med. 2014;20(11):1321–1326.

40. Yamamoto K, Miwa Y, Abe-Suzuki S, et al. Extramedullary hematopoiesis: Elucidating the function of the hematopoietic stem cell niche (Review). Mol Med Rep. 2016;13(1):587–591.

41. Burberry A, Zeng MY, Ding L, et al. Infection mobilizes hematopoietic stem cells through cooperative NOD-like receptor and Toll-like receptor signaling. Cell Host Microbe. 2014;15(6):779–791.

42. Leuschner F, Rauch PJ, Ueno T, et al. Rapid monocyte kinetics in acute myocardial infarction are sustained by extramedullary monocytopoiesis. J Exp Med. 2012;209(1):123–137.

43. Cheshier SH, Prohaska SS, Weissman IL. The effect of bleeding on hematopoietic stem cell cycling and self-renewal. Stem Cells Dev. 2007;16(5):707–717.

44. McKim DB, Yin W, Wang Y, Cole SW, Godbout JP, Sheridan JF. Social Stress Mobilizes Hematopoietic Stem Cells to Establish Persistent Splenic Myelopoiesis. Cell Rep. 2018;25(9):2552–2562 e2553.

45. Fan N, Lavu S, Hanson CA, Tefferi A. Extramedullary hematopoiesis in the absence of myeloproliferative neoplasm: Mayo Clinic case series of 309 patients. Blood Cancer J. 2018;8(12):119.

46. McCabe A, MacNamara KC. Macrophages: Key regulators of steady-state and demand-adapted hematopoiesis. Exp Hematol. 2016;44(4):213–222.

47. McCabe A, Smith JNP, Costello A, Maloney J, Katikaneni D, MacNamara KC. Hematopoietic stem cell loss and hematopoietic failure in severe aplastic anemia is driven by macrophages and aberrant podoplanin expression. Haematologica. 2018;103(9):1451–1461.

48. Loynes CA, Lee JA, Robertson AL, et al. PGE2 production at sites of tissue injury promotes an anti-inflammatory neutrophil phenotype and determines the outcome of inflammation resolution in vivo. Sci Adv. 2018;4(9):eaar8320.

49. Meziani L, Deutsch E, Mondini M. Macrophages in radiation injury: a new therapeutic target. Oncoimmunology. 2018;7(10):e1494488.

50. Mukherjee S, Sheng W, Michkov A, et al. Prostaglandin E2 inhibits profibrotic function of human pulmonary fibroblasts by disrupting Ca(2+) signaling. Am J Physiol Lung Cell Mol Physiol. 2019;316(5):L810–L821.

51. Rastogi S, Willmes DM, Nassiri M, Babina M, Worm M. PGE2 deficiency predisposes to anaphylaxis by causing mast cell hyper-responsiveness. J Allergy Clin Immunol. 2020.

52. Nigrovic PA, Gray DH, Jones T, et al. Genetic inversion in mast cell-deficient (Wsh) mice interrupts corin and manifests as hematopoietic and cardiac aberrancy. Am J Pathol. 2008;173(6):1693–1701.

53. Tong M, Tai HH. Synergistic induction of the nicotinamide adenine dinucleotide-linked 15-hydroxyprostaglandin dehydrogenase by an androgen and interleukin-6 or forskolin in human prostate cancer cells. Endocrinology. 2004;145(5):2141–2147.

