## Supplemental Figures for "15-PGDH Inhibition Activates the Splenic Niche to Promote Hematopoietic Regeneration"

### SUPPLEMENTAL MATERIAL

Supplemental Figures: 4

Supplemental Tables: 0

Supplemental References: 0

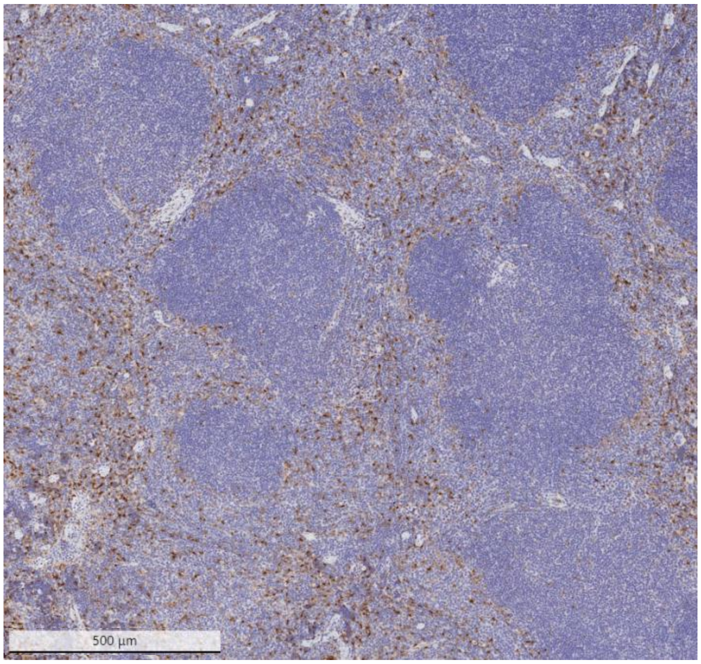


**Supplementary Figure 1. 15-PGDH is expressed in the splenic red pulp.** Representative image of 15-PGDH staining in the murine spleen at 5X magnification. Scale bar represents 500um.

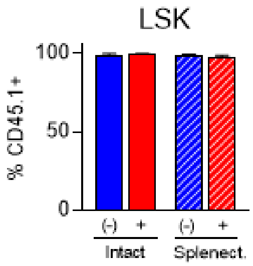


**Supplementary Figure 2. LSKs in splenectomized and intact transplant-recipients are donor-derived.** Quantification of percent CD45.1+ Lineage- Sca1+ c-Kit+ (LSK) cells in the marrow of intact and splenectomized, treated with Veh and PGDHi, 20 days post-transplant. N = 3-6 mice/group.

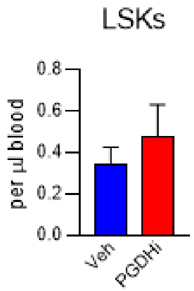


**Supplementary Figure 3. PGDHi does not mobilize HSPCs.** Quantification of Lineage- Sca1+ c-Kit+ cells per ul peripheral blood derived from mice treated 5 days with Veh or PGDHi. N = 15-16 mice/group.

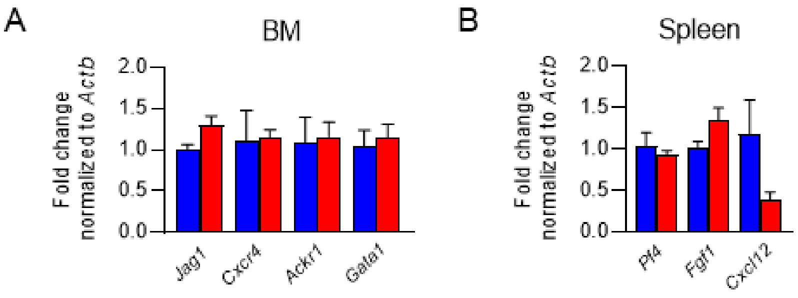


**Supplementary Figure 4. Expression of additional hematopoietic niche-related genes in the BM and spleen of 5 day PGDHi-treated mice.** Relative expression of the indicated genes in BM (**A**) and spleen (**B**). N = 3 mice/group.
